# Non-additive genetic components contribute significantly to population-wide gene expression variation

**DOI:** 10.1101/2023.07.21.550013

**Authors:** Andreas Tsouris, Gauthier Brach, Joseph Schacherer, Jing Hou

**Author notes:** Corresponding authors (J.S.) and (J.H.).

## Abstract

Gene expression variation, an essential step between genomic variation and phenotypic landscape, is collectively controlled by local (*cis*) and distant (*trans*) regulatory changes. Nevertheless, how these regulatory elements differentially influence the heritability of expression traits remains unclear. Here, we bridge this gap by analyzing the transcriptomes of a large diallel panel consisting of 323 unique hybrids originated from genetically divergent yeast isolates. We estimated the broad- and narrow-sense heritability across 5,087 transcript abundance traits and showed that non-additive components account for 36% of the phenotypic variance on average. By comparing allelic expression ratios in the hybrid and the corresponding parental pair, we identified regulatory changes in 25% of all cases, with a majority acting in *trans*. We further showed that *trans-*regulation could underlie coordinated expression variation across highly connected genes, resulting in significantly higher non-additive variance and most likely in some of the missing heritability of gene expression traits.

**Highlights:**

- Diallel panel for dissecting genetic components underlying gene expression variation
- Non-additive components account for 36% of gene expression trait heritability
- Most *cis-* regulatory variation is buffered by additional *trans*-acting variants
- Genes under *trans-* regulation show high non-additive variance and functional coherence

## Introduction

Gene expression is an important molecular step translating genotypes into phenotypes, and its misregulation can have broad consequences on organismal traits^1–4^. Dissecting the regulatory changes that underlie gene expression variation and its heritability can therefore provide important insights into the genetic basis of phenotypic diversity^5–7^. Differences in gene expression between individuals are collectively influenced by variation in local regulatory elements (*cis*-acting) and distant regulatory genes (*trans*-acting)^3^. The interplay between *cis*- and *trans*-acting variants reflect the complex gene regulatory network and underlie the heritable gene expression variation in a population.

The identification of *cis*- and *trans*-regulatory variants in a population typically relies on statistical associations between genetic variants and gene expression levels through large-scale genomic and transcriptomic analyses. Mapping the loci involved in gene expression variation, *i.e.* eQTL (expression Quantitative Trait Loci), often requires a large population to have enough statistical power, especially to detect *trans*-eQTL due to the high number of possible positions to test compared to *cis*-eQTL^8, 9^. As a result, most human eQTL analyses using GWAS are limited to detecting only *cis*-acting variants^10–12^. By contrast, population-wide transcriptomic surveys in model systems consistently show that *trans*-eQTL are more common than *cis*-eQTL and collectively explain a larger fraction of gene expression variance^13–16^.

However, even considering the effects of both *cis*- and *trans*-eQTL, the total phenotypic variance explained remained modest^8, 14, 17–20^. Such variant-centric strategy overlooks the complex interaction between *cis*- and *trans*-regulatory variants acting on the same trait and possibly leads to some extent the observed missing heritability.

Another way to identify *cis*- and *trans-*regulatory variation is through comparative analyses of allele-specific expression (ASE) patterns across pure-bred parental lines and their F1 hybrid^3^. While differences in allelic expression levels both within the hybrid and between parental lines indicate a *cis*-regulatory change, a *trans*-regulatory change will result in different expression levels in the parents but no difference in allelic expression within the hybrid as the *trans*-acting variant act equally on both alleles. Compared to eQTL analyses, ASE-based strategy focuses on the regulatory patterns at the gene level, better captures the biological reality, and is not affected by the statistical challenges like GWAS. However, such a strategy is often limited to one or a few parent-hybrid trios, where population-level variation is ignored.

So far, ASE-based *cis*- and *trans*-regulatory studies have been mainly concentrated in identifying expression divergence between species of *Drosophila*, mice, yeasts and several plants^21–26^. Several studies have used intraspecific pairs, but their scope is limited and they do not reflect the population-wide diversity^27^.

A gene-centric view of the regulatory variation in a population should provide deeper insights into how *cis* and *trans* effects contribute to the heritability of gene expression. From this perspective, the ASE-based method has a unique advantage as heritability can be inferred based on parent-offspring regression, and both additive and non-additive components can be estimated with sufficiently large number of parent-hybrid trios. In comparison, variant-centric methods such as eQTL analysis mainly focus on the additive effect (narrow-sense heritability or *h*^2^), whereas the total genetic effect on phenotypic variance (broad-sense heritability or *H*^2^) remains elusive. Narrow-sense heritability estimated for gene expression traits in humans and other model systems are low, ranges from ∼0.06 to ∼0.30 depending on the studies^8, 14, 17–19^, and evidences based on familial data in human suggest non-additive genetic component could explain part of the missing heritability in gene expression^17, 28^. In fact, gene-centric analysis of regulatory variation at the population scale is still lacking and therefore how regulatory changes contribute to additive and non-additive genetic components in gene expression have never been explored.

Here, we bridge these gaps by generating high quality transcriptomes across a large diallel panel consisting of 323 unique F1 hybrids, originated from 26 genetically divergent parental yeast isolates.

Taking advantage of this diallel design, we estimated broad- and narrow-sense heritability on 5,087 transcript abundance traits. We showed that non-additive variance plays a major role in gene expression variation and accounts for 36% of the phenotypic variance on average. We calculated allele-specific read counts in parent-hybrid trios and characterized the regulatory patterns of approximately 300,000 gene-trio combinations. We found that *trans*-regulatory changes underlie the majority of gene expression variation in the population, with most *cis*-regulatory variation also being exaggerated or attenuated by additional *trans* effects. We further showed that *trans-*regulatory variation is the main force driving the non-additive variance of gene expression traits.

## Results

### The transcription landscape across a diallel hybrid panel

We selected 26 genetically diverse natural isolates from the 1,011 yeast collection as the parental lines to generate a diallel hybrid panel^29^ (Figure 1A). The selected isolates were originated from a wide range of environments (Figure 1B) and different geographical locations (Table S1) to broadly represent de genomic diversity in the species. The nucleotide divergence between each pair of parental lines ranges from 0.03% to 1.10%, with a mean divergence of 0.59% (Figure 1C). Stable haploids of the 26 parental lines were previously generated by replacing the *HO* locus with an antibiotic resistance cassette^30^. We performed pairwise crosses and generated 351 genetically unique hybrids, including 325 heterozygous hybrids and 26 homozygous diploid parental types (Figure 1D, Table S2). We carried out RNA sequencing and obtained high quality transcriptomes for 323 unique hybrids, including all 26 homozygous parental lines, with a mean of 3.7 million reads per sample. Using the previously established pangenome annotations^13, 29^, we obtained the expression levels (as transcript per million or tpm) for 6,186 genes that are expressed in at least half of the samples (tpm > 0), consisting of 5,770 core genes that are invariably present in all 26 parental lines and 422 accessory genes, the majority of which (291/422) correspond to *S. paradoxus* introgressed alleles (Figure 1E, Table S3, Datafile 1). We performed additional RNA sequencing for a subset of biological replicates, specifically for six heterozygous hybrids in duplicates and one parental diploid in triplicates (Table S2). Gene expression levels (tpm) correlate well between replicates, with correlation coefficients ranging from 0.92 to 0.99 with an average of 0.96 (Pearson’s R), indicating high reproducibility in our data (Figure S1A-B).

**Figure 1.**
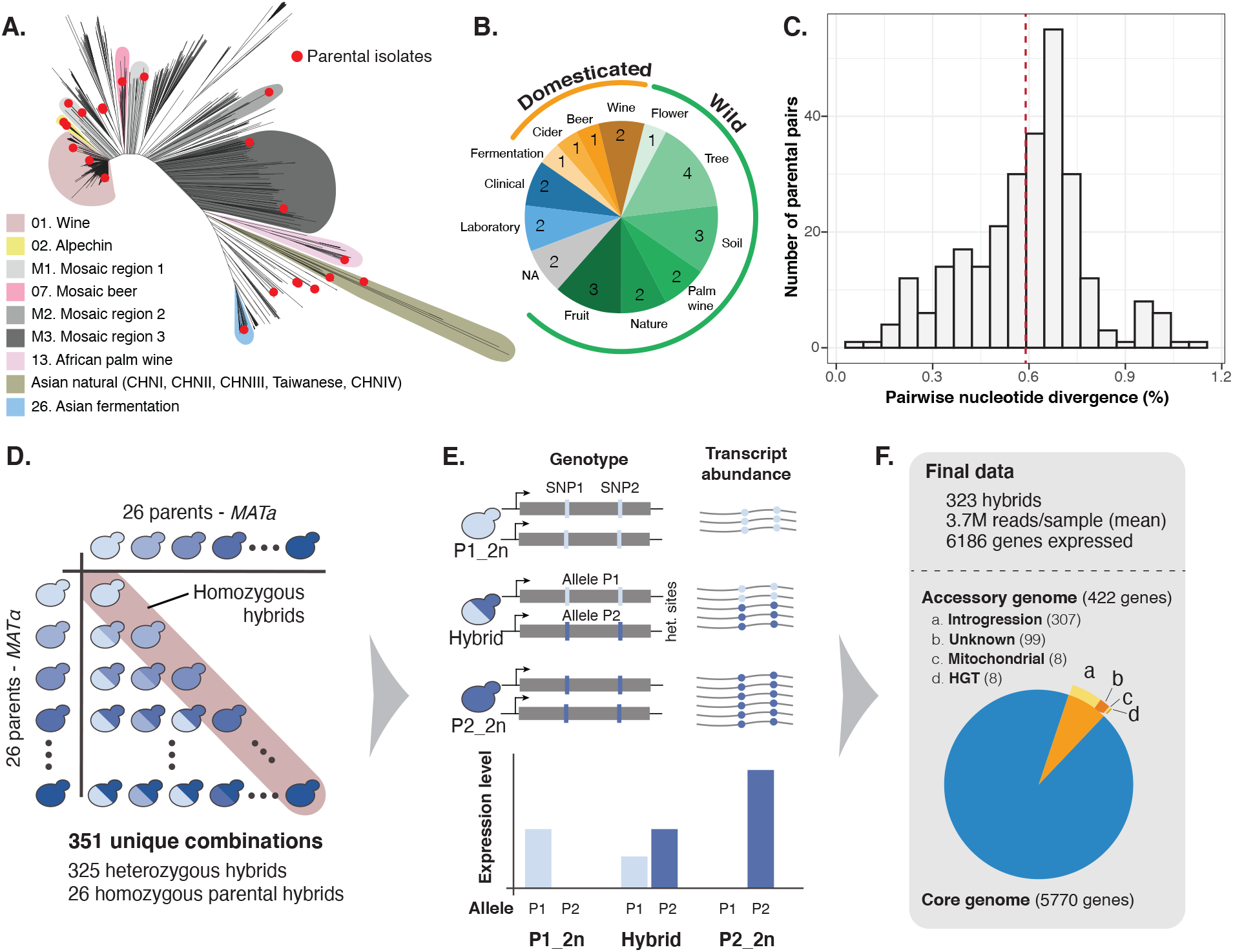
Overview of the diallel design and the transcriptomic dataset. **A.** Neighbor-joining tree based on the genetic diversity of 1,011 yeast isolates^29^. 26 parental isolates selected for the diallel panel are highlighted in red. Representative clades are annotated as in^29^. See Table S1 for detailed origins of the parental isolates. **B.** The ecological origin and distribution of the selected 26 parental isolates. **C.** Pairwise nucleotide diversity among the parental isolates. Mean divergence is indicated in red dashed line. **D.** Schematics of the diallel crossing design. Homozygous parental diploids are highlight in red. See Table S2 for detailed information for the generated hybrids. **E.** Schematics of the data acquisition strategy. For each hybrid the transcript abundance (transcripts per million or tpm) are measured for each annotated ORF, as well as allele specific read counts across all discriminating sites in a given hybrid. Parental allele counts were extracted from the coverage data at the same sites. **F.** Final data metrics and numbers of accessory and core genes included for the subsequent analyses. See Table S3 for detailed annotations.

We previously generated a species-wide pan-transcriptomic dataset involving 969 natural isolates from the same strain collection^13^. To evaluate the general gene expression behavior across the diallel hybrid panel against the previous population-level data, we calculated the mean expression level (*i.e.* abundance) and the mean absolute deviation across samples (*i.e.* dispersion) for each of the 6,186 genes in the final dataset. Both metrics showed good agreements between the diallel panel and the natural population, with a correlation coefficient of 0.79 for abundance and 0.72 for dispersion (Pearson’s R) (Figure S1C-D). These observations suggest that the diallel panel broadly captures the global gene expression variability of the population.

### Non-additive genetic components contribute significantly to gene expression variation

Taking advantage of the diallel design, we calculated the broad- (*H*^2^) and narrow-sense heritability (*h^2^*) for each expression trait by estimating the combining abilities using the Griffing’s model^31^ (Methods). Briefly, for each heterozygous hybrid, the total expression level for a given gene can be decomposed into the average contributions of the parental lines (General Combining Ability or GCA), the contribution due to the combination of the parents in a hybrid (Specific Combining Ability or SCA), and the residual variation that is unrelated to the parental origins. In this context, the additive variance component for a given trait corresponds to the fraction of phenotypic variance explained by the sum of GCA variance from the parents, whereas the non-additive component corresponds to the fraction of phenotypic variance due to the SCA variance. The broad-sense heritability (*H^2^*) is therefore calculated as the sum of additive and non-additive genetic components, and the narrow-sense heritability (*h*^2^) correspond to the additive component only. We obtained *H*^2^ and *h*^2^ estimates for 5,087 out of the 6,186 expression traits (Methods). Across these traits, the estimated *H*^2^ ranges from 0.08 to 0.99 with a median of 0.75, while the *h^2^* ranges from 0.08 to 0.98 with a median of 0.31 (Figure 2A, Table S4). We applied an orthogonal strategy to estimate the *h^2^* based on the genome-wide variants and the kinship matrix across all hybrids using a generalized linear model (Methods). The resulting genome-wide additive heritability (*h^2^g*) is highly correlated with the *h^2^* obtained based on the diallel model (Pearson’s *R*=0.85, P-value < 2.2e-16) (Figure S2A).

**Figure 2.**
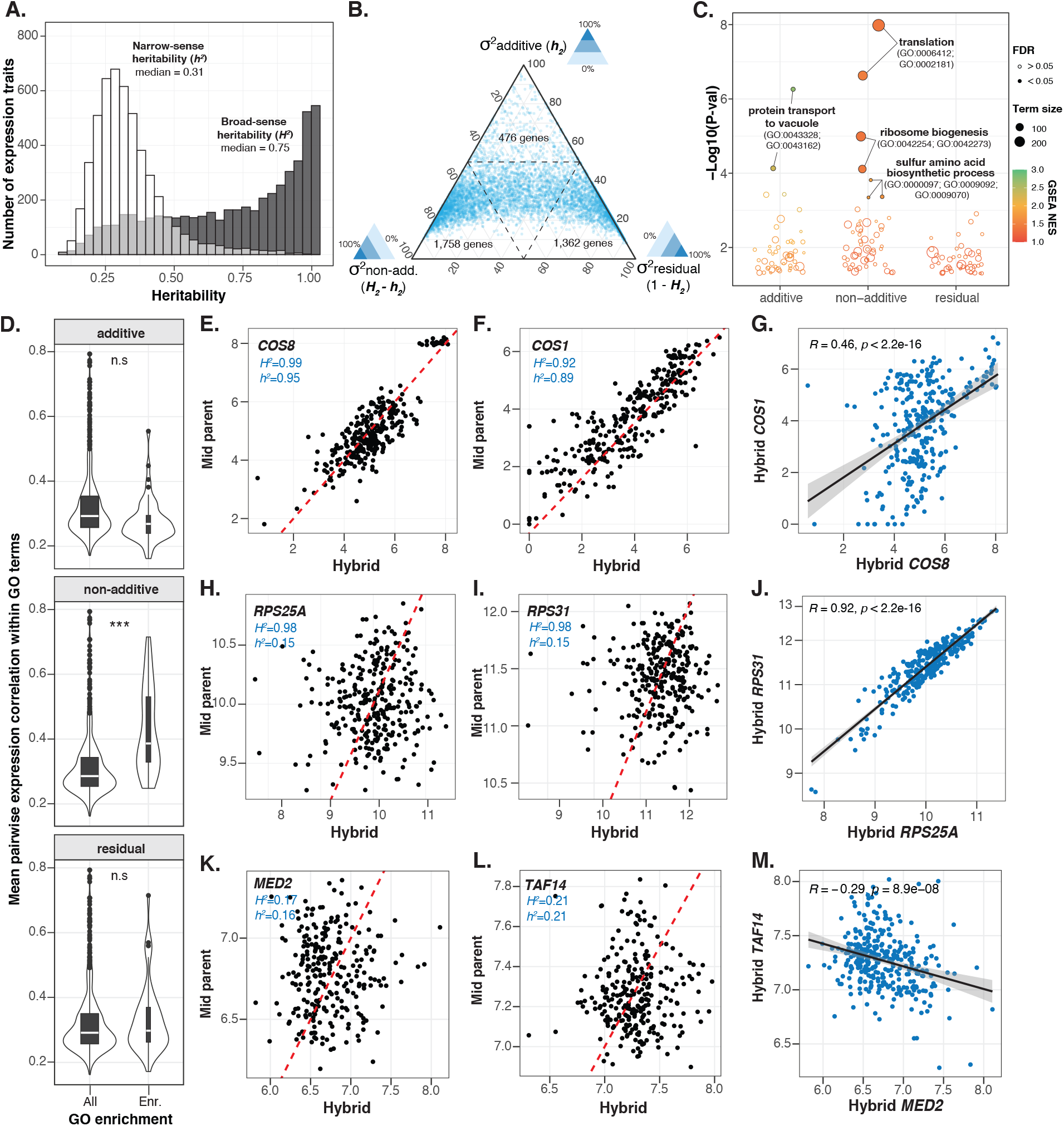
Broad- and narrow-sense heritability for genome-wide gene expression traits. A. Distributions of broad- (grey) and narrow-sense heritability (white) estimates based on the diallel hybrid panel for 5,087 gene expression traits. transcript abundance in white and gray respectively. See Table S4 for all heritability estimates. **B.** Ternary plot showing the percentage of phenotypic variance controlled by additive, non-additive and residual variance components. Dashed line marks the 50% threshold for each dimension. **C.** Gene-set enrichment analysis (GSEA) for additive, non-additive and residual variance components, using standard GO biological process (BP) terms. All terms with a nominal P-value < 0.05 are shown. Color scale corresponds to the normalized enrichment scores. Size of the circles indicate the number of genes annotated on each term. Full circle represents GO terms with an FDR<0.05. See Table S5 for detailed enrichment results. **D.** Mean pairwise expression profile correlation for genes in the same GO term with a nominal enrichment P-value < 0.05 for additive, non-additive and residual variance components. Significant differences between enriched and all terms are indicated with stars (n.s: non­significant; ***: Wilcoxon test P-value < 2.2e-16. **E-M.** Example hybrid-midparent and hybrid profile correlations for genes with high additive (**E-G**), non-additive (**H-J**) and residual (**K-M**) variance components. Example genes correspond to the top two leading edges in the GSEA results. Dashed red lines indicate the one-to-one correlation line (slope = 1) for visual guide. The heritability estimated are indicated for each example gene pairs.

The distribution of different variance components is skewed toward the non-additive component (*H^2^-h^2^*) (Figure 2B). Non-additive variance component accounts for 36% of phenotypic variance on average, with approximately 1/3 of the genome (1,758 genes) mainly under non-additive control (*H^2^- h^2^* > 0 .5). By contrast, only 476 genes are mainly additive (*h^2^* > 0.5). Genes that are mainly controlled by the residual variance (1,352 genes, 1- *H^2^* > 0.5) are characterized by an overall low phenotypic variance across the population (Figure S2B), suggesting variation due to expression stochasticity and random noise.

To explore the relationship between different variance components and gene functions, we ranked genes based on their additive, non-additive and residual variance and performed gene-set enrichment analyses (GSEA) using standard gene ontology (GO) biological process terms. Genes that are highly additive are only significantly enriched for a couple of small terms related to protein transport to vacuole (GO:0043328; GO:0043162) (Figure 2C, Table S5). Highly non-additive genes showed the most significant enrichments, specifically for terms related to translation (GO:0006421; GO:0002181), ribosomal biogenesis (GO:0042254; GO:0042273) and sulfur amino acid biosynthesis (GO:0000097; GO:0009092; GO:0009070) (Figure 2C, Table S5). No significant enrichment (FDR < 5%) were found for genes that show high residual variance (Figure 2C, Table S5). These results suggest that genes with similar biological functions could show similar variance component profiles, most notably for genes that are mainly controlled by non-additive variance.

We examined the expression coherence (*i.e.* co-regulated expression patterns) by calculating the pairwise expression profile similarity among genes that belonged to all GO terms with a nominal enrichment P-value < 0.05 for additive, non-additive and residual variance components (Figure 2D). Compared to all GO terms, terms enriched for non-additive variance showed significantly higher pairwise gene expression profile similarity on average (one-sided Wilcoxon test, P-value < 2.2e-16), contrasting to terms that are enriched for additive or residual variance (Figure 2D). As examples, we took the top two leading edges for GO terms with the lowest enrichment P-value for each of the three variance component rankings (Figure 2E-M). The top leading edges that are highly additive correspond to *COS8* (*H^2^*=0.99, *h^2^*=0.95) and *COS1* (*H^2^*=0.92, *h^2^*=0.89), both are ubiquitin cargos involved in protein transport to vacuole via the multivesicular body sorting pathway (GO:0043328). Both genes are indeed showing high additive effect as evidenced by the correlated expression levels between the hybrids and the mid-parent values (mean expression between the corresponding homozygous parental diploids) (Figure 2E-F). However, these two genes are not co-regulated across the hybrids, with a profile similarity of ∼0.46 between the gene pair (Pearson’s R) (Figure 2G). Top leading-edge genes that are highly non-additive correspond to *RPS25A* and *RPS31,* both are ribosomal proteins involved in translation. Both these genes are characterized by low correlations between the hybrid expression levels and the mid-parent values (Figure 2H-I). Yet, these two genes show highly correlated expression profiles, suggesting co-regulated expressions in the hybrids that is not predicted based on the parental mean. Finally, for genes that show high residual variance, the most significant GO term corresponds to RNA polymerase II transcriptional preinitiation complex assembly (GO:0051123), with a nominal enrichment P-value = 0.001 and an FDR = 0.76. The top leading edges correspond to *MED2*, a subunit of the RNA polymerase II mediator complex; and *TAF14,* a DNA binding protein involved in RNA polymerase II transcription initiation and in chromatin modification. As expected, no correlation is observed between the hybrid expression level and the mid-parent value (Figure 2K-L), nor between the expression profiles across hybrids (Figure 2M).

Overall, the diallel design allowed us to effectively decompose the variance components associated with the majority of gene expression traits. Non-additive genetic variance contributes significantly to gene expression variation, explaining 36% of the phenotypic variance on average. Different variance components contribute unequally to genes with different cellular functions and the non-additive component showing the highest functional coherence in the hybrid panel.

### Widespread transcriptomic buffering via *cis*-*trans* compensation

The genetic component of gene expression variation can be attributed to regulatory variants acting in *cis* and/or in *trans*. In principle, local DNA sequence variation that impact gene expression (*e.g.* mutations in promoter regions) are *cis*-acting, whereas *trans*-regulatory variants act distantly (*e.g.* transcription factors) and can occur anywhere in the genome. The diallel panel consists of pairwise combinations of a large number of parent-hybrid trios. For each trio, the regulatory variation in *cis* or *trans* can be determined by comparing the allelic expression in the hybrid to the corresponding expression levels in the parental lines. Specifically, for a given gene, if the expression difference between the two parental lines is due to *cis*­regulatory change, the corresponding alleles will result in allele-specific expression (ASE) in the hybrid (Figure 3A). Conversely, in the case of *trans-regulatory* change, no allele-specific expression would be observed as the *trans*-acting factor impact equally both alleles in the hybrid background (Figure 3B).

**Figure 3.**
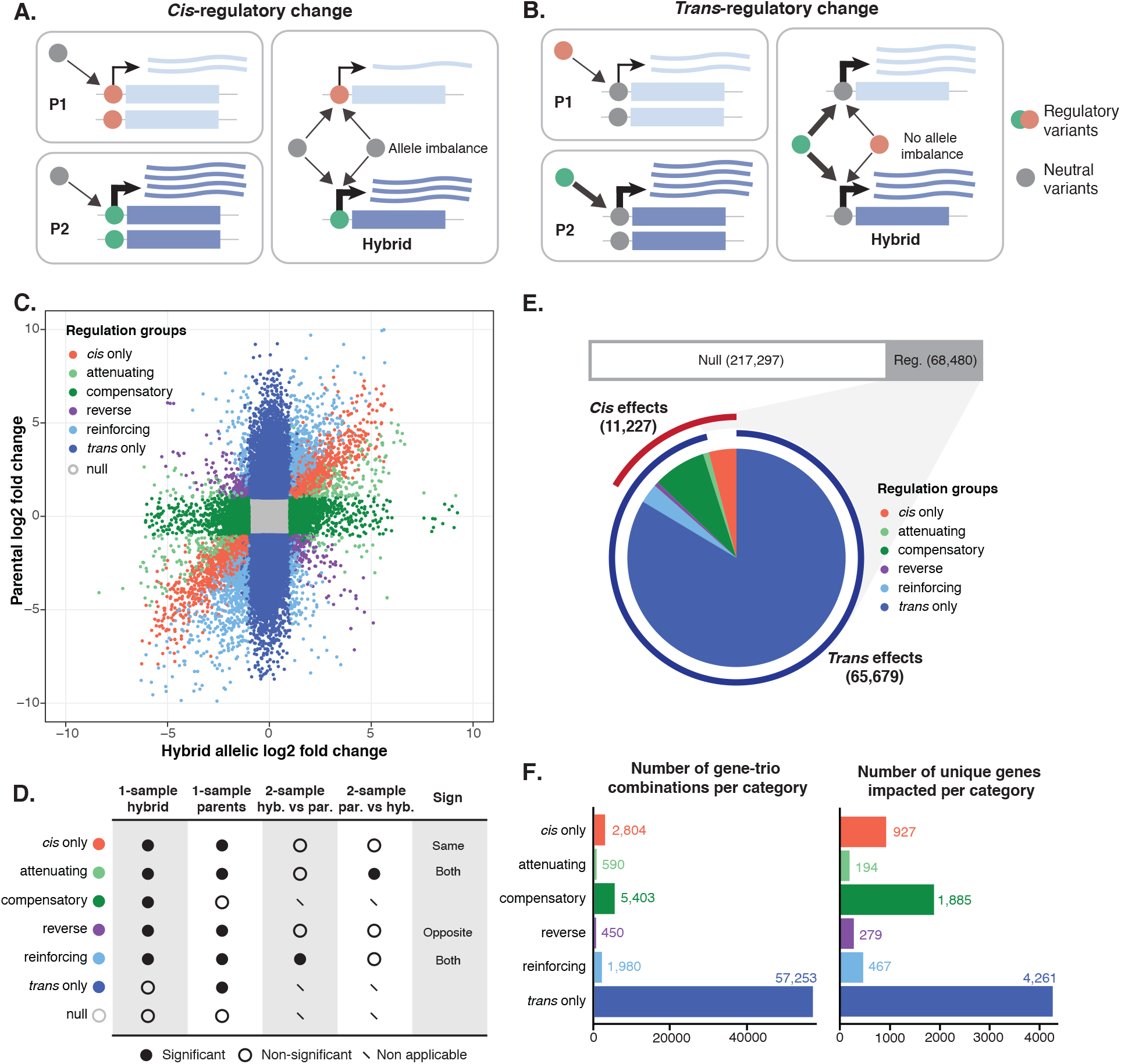
Systematic characterization of regulatory variation across the population. A-B. Schematic depiction for *cis*- (**A**) and *trans*- (**B**) regulatory variation and the resulting allele-specific expression patterns across parent-hybrid trios. **C.** Regulatory variation patterns identified across 285,777 gene-trio combinations. Log2 fold changes between alleles in the hybrid and between the parental lines at the same sites are indicated on x- and y-axis, respectively. Different regulatory patterns are color coded. **D.** Criteria for classifying different regulatory patterns based on 1- and 2-sample ASE test significance. See detailed description in Methods. **E.** Number and distribution for different regulatory patterns across the whole dataset. Upper bar indicates the number of the “null” category vs. other categories with significant regulatory changes. Pie chart indicate the proportions of all significant regulatory patterns, with outer ring indicating cases that are under *cis*- (red) or *trans*- (blue) controls. **F.** Number of gene-trio combinations per regulatory category (left panel) and the number of unique genes impacted in the corresponding groups (right panel). See detailed results in Datafile 2.

We determined the allele-specific read counts at each discriminating site within gene open reading frame (ORF) in the heterozygous hybrids and in the corresponding parental lines. We removed low coverage sites (sum of hybrid allele counts < 30 and sum of parental counts < 60) and excluded cases where one of the gene copies is absent in either one or both parents. We also excluded any trios that showed inconsistent chromosome-level allele balance patterns in the hybrid or between the parental lines (Methods). In total, 179 unique parent-hybrid trios were retained for further analyses, comprising ∼1.2 M sites across 285,777 gene-trio combinations (Datafile 2). The final data covered 5,089 genes, including 219 accessory genes from *5. paradoxus* introgression (Datafile 2). On average, ∼1,600 genes contained at least one such discriminating site per trio (Figure S3A-B), for which the regulatory variation can be inferred by comparing the allelic ratio change in the hybrid to the parental ratio at the same sites.

For each of the 285,777 gene-trio combinations, we performed 1-sample ASE tests both in the hybrid and between the parental pair by considering the allele counts across all sites within the same gene (Methods). For cases that showed significant allelic ratio differences (|log2 fold-change| > 1 & FDR < 0.05) in both the hybrid and the parents, 2-sample ASE tests were subsequently performed to identify significant changes between the hybrid and parental allelic ratios (FDR < 0.05) (Methods). Based on test results, we categorized cases into different regulatory patterns (Figure 3C-D). Overall, ∼76% (217,297out of 285,777) of all cases showed no significant allelic expression differences in either the hybrid or the parents and are classified as the “null” type, while ∼24% (68,480 out of 285,777) displayed significant regulatory variation (Figure 3E). Among these 68,480 cases, ∼16% (11,227 out of 68,480) showed evidence for *cis* effect while ∼96% (65,679 out of 68,480) showed *trans* effect, with ∼12% (8,423 out of 68,480) showing significance for both (Figure 3E). Cases that are exclusively controlled by *cis* or *trans* effects represent ∼4% (2,804 out of 68,480) and ∼84% (57,253 out of 68,480), respectively (Figure 3F). Cases with combined *cis* and *trans* effects were further grouped into four distinct regulatory patterns (Figure 3C-D). In the “attenuating” group (∼0.9%, 590 out of 68,480), the *cis* effect in the hybrid is decreased in magnitude compared to the parental expression levels by additional *trans* factors (Figure 3D). The “reinforcing” group (∼3%, 1,980 out of68,480) describes the opposite event where the *cis* effect in the hybrid is exaggerated in magnitude compared to the parental expression variation (Figure 3D). The “compensatory” group (∼8%, 5,403 out of 68,480) corresponds to the majority of cases with both *cis* and *trans* effects. In these cases, the *cis* regulatory effect is completely cancelled out by additional *trans* factors, resulting in significant allele-specific expression within the hybrid but no expression difference between the parents (Figure 3D). Finally, the “reverse” group corresponds to extreme *cis*/*trans* interaction events, where the allelic variation in the hybrid is in complete opposite direction than the parental variation (Figure 3D). These events are rare and represent 0.7% (450 out of 68,480) of all cases with significant regulatory variation (Figure 3D-F).

Globally, *trans*-regulatory variation is more common than *cis*, which is consistent with previous observations using eQTL analyses^8,13^. Remarkably, the majority of *cis*-regulatory variation are influenced by additional *trans* effect. Among all gene-trio combinations showing *cis*-regulatory variation, ∼75% (8,423 out of 11,227) also show *trans*-regulatory changes and most of such *trans* effects act in opposite direction relative to the observed *cis* effect, specifically in the “attenuating” (590 out of 11,227; ∼5%) and the “compensatory” (5,403 out of 11,227; ∼48%) groups. These observations suggest that *cis*-regulatory variations are globally compensated in *trans*, resulting in a general buffering effect at the transcriptome level.

### Regulatory variation in *trans* plays a major role in non-additive heritability

In our diallel scheme, the regulatory variation associated with each gene is characterized across multiple hybrid-parent trios, which allows us to probe how different regulatory changes at the individual level (*i.e.* any given gene-trio combination) impact global gene expression features such as heritability and different variance components across the full hybrid panel. In total, 3,791 genes (out of 5,089 with discriminating sites) are characterized in at least 10 hybrids, which were further analyzed (Figure S4A, Datafile 2). For a given gene, the regulatory variation across different hybrids can be relatively conserved. For example, *GTO1,* encoding for a glutathione transferase, is most exclusively controlled in *cis* (Figure 4A). The *cis­*regulatory change is due to a single parental line, Y12, an isolate associated with Asian fermentation where *GTO1* is known to be differentially overexpressed compared to other subpopulations^13^. Another example with conserved regulatory variation is seen for *CYC7*, encoding for an isoform of cytochrome c, in which case the regulatory variation among different hybrids is mainly *trans-acting* (Figure 4B). For cases with combined *cis* and *trans* effects such as the “compensatory” pattern, the regulatory variation across hybrids can be complex, such as the case for *RPL4B,* encoding for a ribosomal subunit (Figure 4C). In this case, while the majority of hybrids show a “compensatory” pattern, other patterns are also seen due to different combinations of *cis-* and *trans-* acting factors in different hybrids.

**Figure 4.**
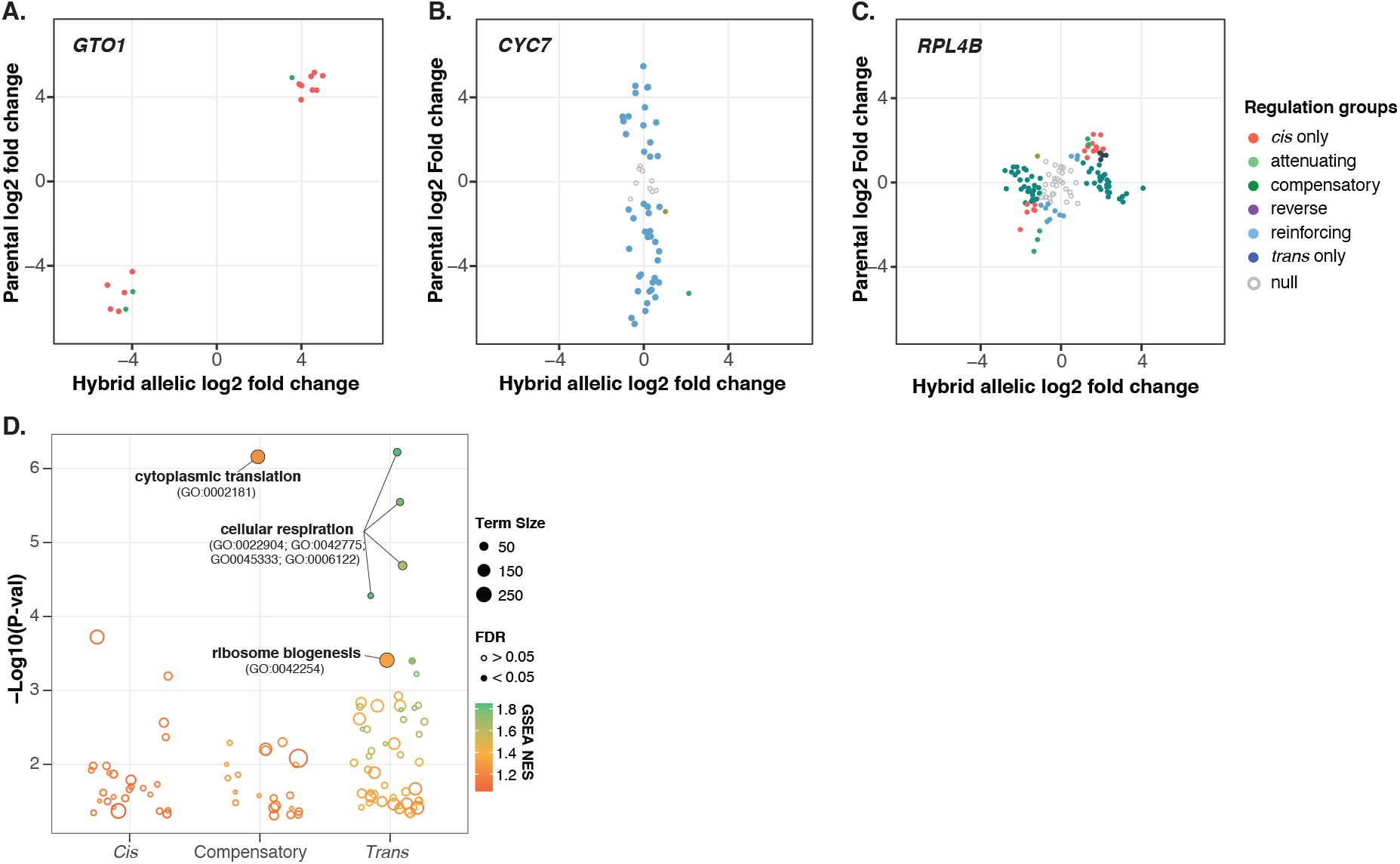
Functional enrichments for genes with distinct regulatory patterns and associated examples. A-C. Examples of genes that are mainly under *cis* (**A**), *trans* (**B**) and compensatory (**C**) regulatory controls. Log2 fold changes between alleles in the hybrid and between the parental lines at the same sites are shown. Different regulatory patterns are color coded. **D.** GSEA results for genes that are mainly under *cis*, *trans* and compensatory regulatory controls. All go terms with a nominal P-value < 0.05 are shown. Terms with significant enrichment (FDR < 0.05) are shown as solid circles. Color scale correspond to the normalized enrichment scores. Circle sizes indicate the number of genes associated with each term. See detailed enrichment results in Table S6.

We ranked genes based on the number of “*cis* only”, “*trans* only” and “compensatory” cases across the hybrid panel and performed GSEA based on standard GO biological processes terms in order to see if genes with similar cellular functions could display conserved regulatory patterns across the population. Significant enrichments were observed for the “compensatory” and the *“trans* only” groups (Figure 4D). Genes that showed the most “compensatory” patterns are enriched for cytoplasmic translation (GO:0002181) (Figure 4D, Table S5). On the other hand, genes that showed the most *“trans* only” cases are enriched for cellular respiration related processes (GO:0022904; GO:0042775; GO:0045333; GO:0006122), glycogen metabolic processes (GO:0005977; GO:0032787) and ribosome biogenesis (GO:0042254) (Figure 4D, Table S5).

We further categorized genes based on the major regulatory pattern present across hybrids (Methods). We examined the behavior of genes across various global expression features (Figure 5A-C) and genetic variance components (Figure 5D-F). As expected, genes that are mainly under *cis* control show overall low expression abundance (Figure 5A), high expression dispersion (Figure 5B) and low connectivity (Figure 5C), all of which are features associated with genes that tend to show the most *cis*-eQTL based on our previous population-level transcriptomic analysis^13^. Indeed, *cis*-regulated genes also show significantly higher proportion of additive variance *(h^2^)* (Figure 5D). By contrast, genes that are mainly under *trans* control are highly expressed (Figure 5A), less dispersed than *cis* controlled genes (Figure 5B) and highly connected on the global co-expression network (Figure 5C). Lastly, genes that show the most compensatory patterns display intermediate features compared to *trans* controlled genes for expression abundance and connectivity (Figure 5A&C). However, this group of genes show the lowest expression dispersion, which is consistent with the *cis-trans* compensation effect (Figure 5B). In general, both compensatory and *trans* controlled genes showed significantly higher proportion of non-additive variance (*H^2^-h^2^*) compared to genes that are mainly *cis-*regulated (Figure 5E). We further compared the hybrid expression abundance to the mid-parent value for each gene-trio combination and identified cases that deviated from the parental expectations across different regulatory variation groups (Figure S4B) (Methods). The majority of cases with expression deviation is attributed to *trans*-regulatory changes (∼46%, 3,166/6,894) (Figure S4C). Nonetheless, the “compensatory” group showed higher proportion of cases with mid-parent deviation as well as significantly higher magnitude of such deviation compared to the “*cis* only” and “*trans* only” groups (Figure S4C-D).

**Figure 5.**
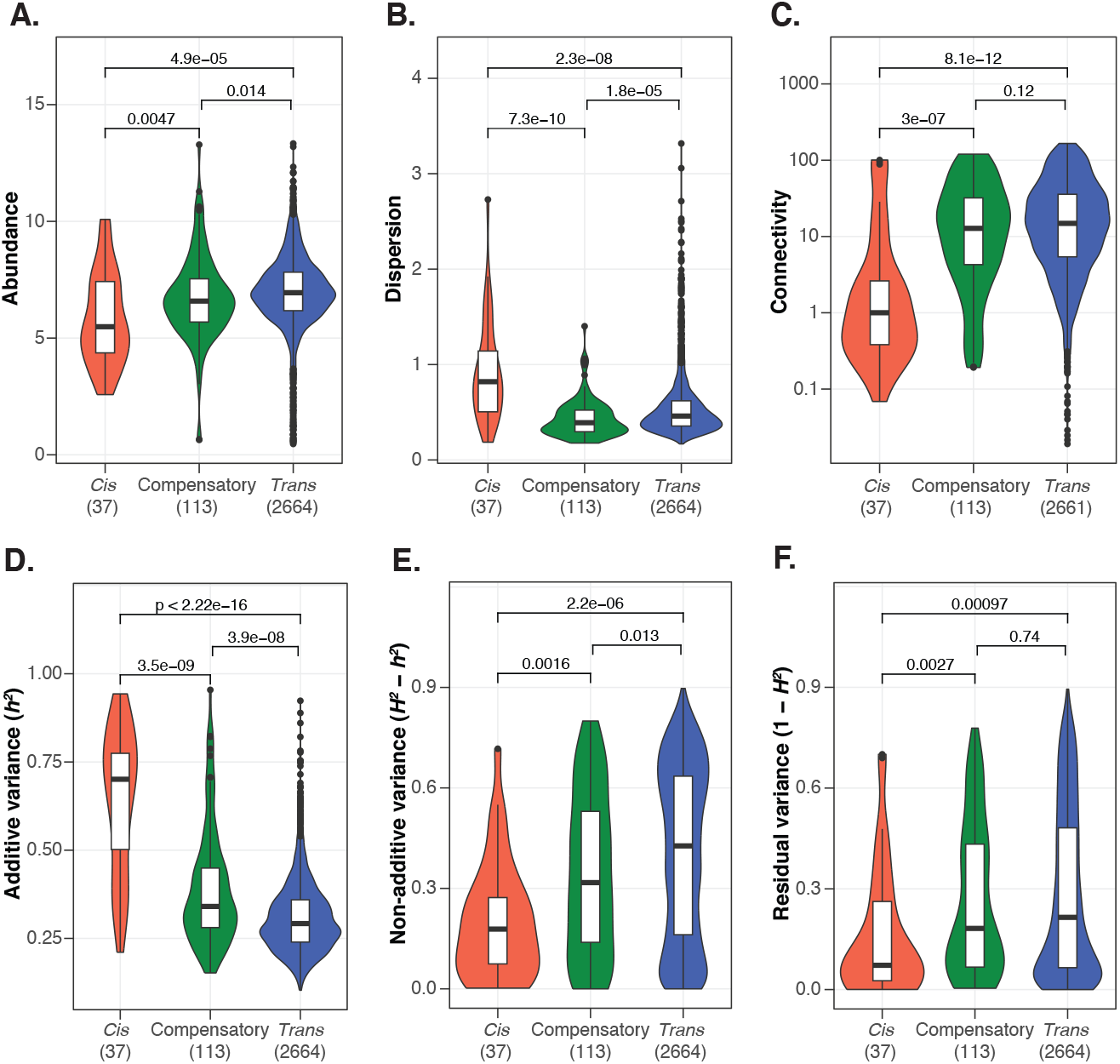
Functional features associated with different regulatory variation patterns. Global gene expression features and their associations with major regulatory patterns (**A-C**). **A.** Expression abundance as the mean log2(tpm+1) across diallel panel. **B.** Expression dispersion calculated as the mean absolute deviation of log2(tpm+1) across samples. **C.** Connectivity calculated as the weighted network connectivity across previous population-level pan-transcriptomic analyses with 969 natural isolates^13^. P-values indicated corresponds to two-sided Wilcoxon tests. Variance components and their associations with major regulatory patterns (**D-F**). **D.** Additive variance component (*h^2^*). **E.** Non-additive variance component (*H^2^-h^2^*). **F.** Residual variance component (1-*H^2^*). The number of genes belonging to different regulation types are indicated on the x-axis. See assigned major regulatory patterns for genes in Table S7.

Overall, our data suggest that *trans*-regulatory changes underlie highly connected, core cellular processes and is the major contributor to the non-additive variance component in gene expression. *Cis-trans* compensation events, while less frequent than *trans*-only cases, contribute significantly to non-additivity as well due to higher magnitude of parent-hybrid expression deviation.

### *S. paradoxus* introgression genes show more *cis*-regulation than their *S. cerevisiae* counterparts

Genes originated from *S. paradoxus* introgression constitute ∼30% of the *S. cerevisiae* accessory genome^29^ and contribute significantly to heritable gene expression variation at the population-wide pan-transcriptome landscape in the species^13^. These introgressed genes are mainly found in specific subpopulations such as the Alpechin clade^29^. In the diallel hybrid panel, two Alpechin isolates were included as parental lines (Table S1), which offered a unique opportunity to examine and compare the regulatory variation among alleles within (*S. cerevisiae* vs *S. cerevisiae*) and between species (*S. cerevisiae* vs *S. paradoxus*). In total, the regulatory variation for 219 unique introgressed genes were included in our dataset, corresponding to 3,192 gene-trio combinations (between species alleles) (Datafile 2). Among the 219 introgressed genes, 202 also showed discriminating sites among their *S. cerevisiae* counterparts, corresponding to 11,871 gene- trio combinations (within species alleles) (Datafile 2). The number of gene-trio combinations with between species allele pairs is mainly associated with the two parental lines from the Alpechin clade, whereas the number of cases with within species allele pairs is more randomly distributed across parental lines (Figure S5A). Globally, for the same set of 202 genes, between species allele pairs show significantly more regulatory variation than within species allele pairs (Figure 6A). Such difference is mainly due to higher proportions of *cis*-regulatory changes observed for the between species allele pairs, including all *cis-trans* interaction patterns (Figure 6A, Figure S5B).

**Figure 6.**
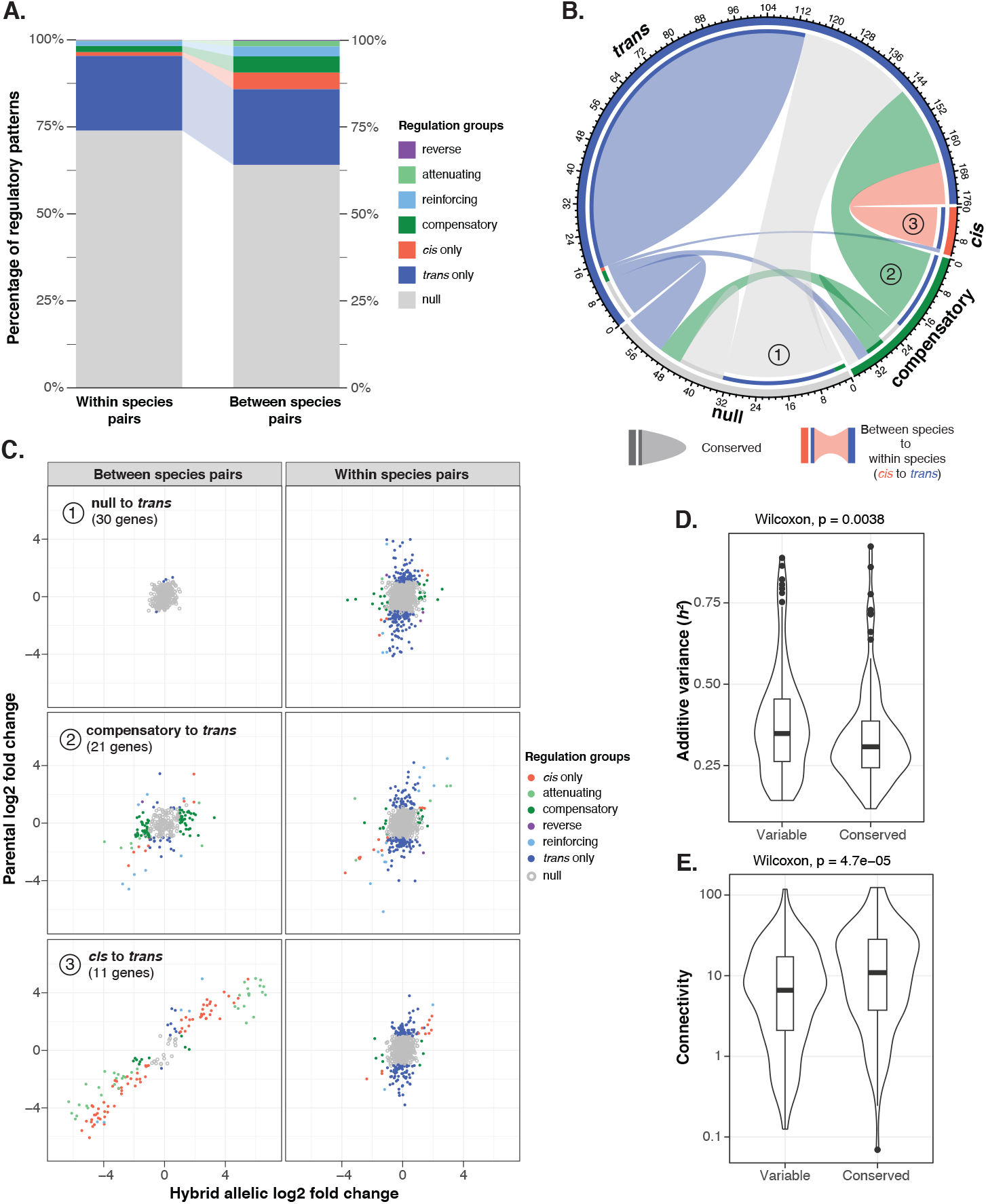
Differential preference of regulatory patterns between introgressed and native alleles A. Distribution and proportions of different regulatory patterns for within and between species allele pairs. **B.** Chord diagram depicting the direction and number of switches in main regulatory modes from between species to within species allele pairs for the same genes. Outer axis indicates the number of gene events. Inner arcs indicate the directional change with the flow starting from between species (with inner arc, color correspond to the destination) to within species (without inner arc). Closed arcs correspond to conserved regulatory pattern. **C.** Major switches observed in (**B**). Regulatory variation for between and within species allele pairs are shown. Log2 fold changes between alleles in the hybrid and between the parental lines at the same sites are indicated on x- and y-axis, respectively. Different regulatory patterns are color coded. **D.** Additive variance component associated with genes that show variable and conserved regulatory patterns for between and within species allele pairs. Wilcoxon test P-value is shown. **E.** Connectivity associated with genes that show variable and conserved regulatory patterns for between and within species allele pairs. Wilcoxon test P-value is shown. Assigned regulatory patterns and types of switches for between and within species alleles are found in Table S8. species to within species allelic regulation modes are “null to *trans*” (30 genes), “compensatory to *trans*” (21 genes) and *“cis* to *trans’”* (11 genes) (Figure 6C). Each of these switches impact the majority of genes within their respective regulation pattern found in the introgressed alleles, specifically with 30 “null to *trans*” out of 46 null in total, 21 “compensatory to *trans*” out of 32 compensatory in total, and 11 “*cis* to *trans*” out of 12 *cis* regulated genes in total (Figure 6B-C). Overall, genes with conserved regulatory patterns between the introgressed and the native alleles tend to show higher expression connectivity and lower additive phenotypic variance *(h^2^)* (Figure 6D-E). These observations suggest that heritable variation of gene expression related to introgression is collectively influenced by the functional integration of such genes to the global expression network (*i.e.* connectivity) as well as their regulatory variation due to interspecies differences between *S. paradoxus* and *S. cerevisiae*.

To understand the global regulatory preferences for introgressed alleles and the native *S. cerevisiae* alleles at the gene level, we assigned the major regulatory pattern observed for between and within species allele pairs and compared the two assignments for the same gene (Methods). In total, 116 out of the 202 genes showed conserved regulatory patterns, the majority (99/116) of these cases are regulated in *trans* both for the introgressed alleles and the *5. cerevisiae* alleles (Figure 6B). The remaining 86 genes showed various types of switches of regulation mode between the introgressed alleles and the *5. cerevisiae* alleles (Figure 6B). Most remarkably, all mainly *cis-*regulated introgressed alleles showed mainly *trans*-regulatory variation for their *5. cerevisiae* counterparts (Figure 6B). For example, the *COX6* gene encodes for the subunit VI of the cytochrome c oxidase and is essential for cellular respiration. The *5. paradoxus* introgressed allele show lower expression levels due to *cis*-regulatory variation, whereas the *5. cerevisiae* alleles are mainly *trans*-regulated (Figure S5C). The most important types of switches going from between

## Discussion

Gene expression variation is a key molecular intermediate in the phenotypic landscape of a species. Understanding how *cis-* and *trans-*regulatory changes differentially influence the heritability of gene expression variation is therefore essential to understand the path from genome to traits. Taking advantage of a large diallel hybrid panel, we estimated the broad- and narrow-sense heritability (*H^2^* and *h^2^*) across genome-wide gene expression traits and showed non-additive components explain higher proportion of the phenotypic variance compared to additive variance only (36% *vs*. 31% on average). We systematically characterized the gene-level regulatory patterns across the population and showed that *trans*-regulatory changes are the main driver underlying the non-additive variance.

It has been previously proposed that *trans*-regulatory changes might have higher tendency to cause gene misexpression than *cis*-variation in the hybrid compared to the parental mean and therefore lead to non-additive variance^32^. However, our data, which tested multiple parental combinations for the same set of genes, show that *trans*-regulatory variation is equally likely to cause hybrid expression deviation from the parental mean than *cis*-variation. While *trans*-regulatory variation appears to be the most important contributor of non-additive variance components in gene expression, the precise underlying mechanism is still unclear.

Our results suggest there are two main paths towards high non-additive variance in gene expression due to *trans* effects. The first one is through *cis*-*trans* interactions, most notably the compensatory effect that completely mask the *cis* effect by additional *trans* factors. Indeed, according to our data, 75% of all *cis* effect are exaggerated or attenuated by additional *trans* variation, and 50% resulted in complete *cis*-*trans* compensation. All *cis*-*trans* interaction patterns show higher tendency for hybrid misexpression and/or higher magnitude of such misexpression. This observation is also supported by previous analysis of the expression of ∼30 genes in a hybrid of two *Drosophila* species^33^. Furthermore, we also showed that genes under *cis-trans* compensation tend to show higher fraction of non-additive variance. In principle, such effect will decrease the power for statistical association in eQTL analyses and will likely result in the missing heritability problem that is commonly observed for gene expression traits.

The second path to non-additive variance is through coordinated expression change in highly connected co-expression modules. Previously, we generated and analyzed the population-level pan-transcriptome across ∼1,000 natural yeast isolates^13^. We showed that the global transcriptome landscape is consistent with a central, highly connected co-expression network and an auxiliary, lowly connected network consists of subpopulation-specific expression signatures^13^. The highly connected co-expression network is robust to genetic variation in the population and showed significantly less eQTL than the auxiliary level^13^. Here, with the diallel hybrid panel, we show that genes with high non-additive variance tend to be highly connected on the global expression network and are often controlled in *trans*. Such non-additive variance is invisible to canonical association tests, which could partly explain the robustness of the co-expression modules to genetic variation in the population. Furthermore, our analyses on the set of 202 introgressed genes also support the link between connectivity and non-additive variance. Introgressed genes that are both *trans*-regulated between interspecies and intraspecies allele pairs show significantly higher connectivity and non-additive variance than cases with variable regulatory patterns.

Overall, our study highlights that the transcriptome is globally buffered at the genetic level. Highly connected co-expression modules are robust to genetic variation in the population, either through *trans* compensation of *cis*-regulatory variation, or through coordinated *trans-*regulatory changes alone, with possible purging of *cis*-effect variants. Such buffering effect could result in significant non-additive variance that is not detectable via genome-wide association surveys and contribute to missing heritability.

## Materials and methods

### Description of the parental strains and diallel scheme

A genetically diverse set of 26 parental strains was selected from the 1,011 strains collection with the focus of capturing as much of the genetic, ecological and geographical diversity of the species as possible (Table S1). Two stable haploid strains, *MAT*a and *MAT*alpha, carrying the *KanMX* or a *NatMX* cassettes in the *HO* locus respectively, were established for each parent, giving a total of 52 strains^30^. Parental strains of opposite mating types were crossed by overlaying haploid cells in a matrix on YPD media (1% yeast extract, 2% peptone and 2% glucose) and incubating them at 30°C overnight. The cells were then transferred to YPD media with G418 (200 mg/ml) and nourseothricin (100 mg/ml) and incubated at 30°C overnight to select for hybrid cells. We then transferred the selected hybrids on YPD media with nourseothricin and G418 to remove any remaining haploid cells. All procedures were done using the replicating robot ROTOR (Singer Instruments). In total, we obtained 351 genetically unique hybrids.

We quantified the growth rates of each hybrid in liquid synthetic complete (SC) media with 2% glucose for 48 hours (initial OD_600nm_: 0.1) using a 96-well microplate reader (Tecan Infinite F200 Pro). The hybrids were then grown in 96-format deepwell plates until mid-log phase (OD_600nm_ ∼0.3). A 750 μL suspension of each sample was then transferred to a 96-well filter plate (Norgen, #40008) where the media was eliminated by applying vacuum (VWR, #16003-836). Immediately after the media was eliminated, we flash-froze the cells in liquid nitrogen and stored them at -80°C.

### Sample preparation

We extracted the mRNA from the hybrids using the Dynabeads® mRNA Direct Kit (ThermoFisher #61012). Cells were lysed with glass beads and incubated at 65°C for 2 minutes and mRNA was then selected with two rounds of hybridization of their polyA tails to magnetic beads coupled to oligo(dT) residues.

We prepared cDNA sequencing libraries using the NEBNext® Ultra™ II Directional RNA Library Prep Kit (NEB, #E7765L) and following the manufacturer’s protocol. Briefly, 5 μL of purified mRNA is fragmented with a 15-minute incubation at 94°C and then reverse transcribed to cDNA. Next, a NEBNext Adaptor is ligated to the cDNA and finally a unique combination of dual indexes (manufactured by IDT®) is added to allow multiplexed sequencing. Finally, barcoded cDNA is amplified with 9 rounds of PCR (denaturation 10 seconds at 98°C, annealing/extension 75 seconds at 65°C). The amplified and barcoded cDNA fragments were eluted in 15 μL of 0.1X TE.

We quantified the concentration of the cDNA libraries with the Qubit ™ dsDNA HS Assay Kit (Invitrogen ™) in a 96-well plate using a microplate reader (Tecan Infinite F200 Pro) with an excitation frequency of 485 nm and emission of 528 nm. We pooled 1 μL of each library and fragment size was assessed with Bioanalyzer 2100 (Agilent™) using the High sensitivity DNA kit (#5067-4626). Finally, we generated equimolar sequencing pools of 96 samples.

The pools were sequenced for 75 bp single-end with Nextseq 550 (Illumina™) sequencer at the EMBL Genomics Core Facility. After demultiplexing, we obtained 3.7 million reads per sample on average. The number of reads obtained in each sample are found in Table S2.

### Quantification of mRNA abundance

The raw reads of each sample were mapped to a custom reference genome using STAR^34^ with the following parameters:

--outSAMtype BAM SortedByCoordinate \

--outFilterType BySJout \

--outFilterMultimapNmax 20 \

--outFilterMismatchNmax 4 \

--alignIntronMin 20 \

--alignIntronMax 2000 \

--alignSJoverhangMin 8 \

--alignSJDBoverhangMin 1

We generated a custom reference genome containing the 16 chromosomes of the *S. cerevisiae* reference genome (R64_nucl) and all the accessory ORFs (n=665) present in the parental strains as defined by the 1,011 yeast genomes project (Peter *et al*., 2018) as additional chromosomes. In total, 323 hybrids had sufficient reads and were used in the subsequent analyses. We counted the reads aligning to each gene of the reference (n=6,285) and accessory (n=665) genomes with the featureCounts function of the R package subread^35^ with the *countMultiMappingReads=F* parameter to eliminate multi-mapped reads. For a given hybrid, if accessory genes that have orthologs in the reference genome were annotated as absent, we merged their reads counts to those of their reference genome counterparts. We then normalized mRNA abundance to the gene level by calculating the transcripts per million (tpm) of each gene. This gave us a list of tpm values for 6917 genes. We filtered out genes that have a zero tpm value in more than half of all samples, leading to a final dataset of 6186 genes. Raw counts and tpm values can be found in Datafile 1.

### Calculating gene-level expression features

We calculated the overall expression features for abundance, dispersion and connectivity for each gene across the diallel and across 969 natural isolates where the transcriptomes were characterized previously^13^. The abundance and dispersion are calculated as previously described^13^. Briefly, the expression abundance was calculated as the mean tpm levels across samples where the gene is present based on genome annotations. The dispersion corresponds to the mean absolute deviation of tpm levels across samples. The connectivity for a given gene is defined as the weighted network connectivity across all expressed genes. The connectivity is calculated using the softConnectivity function in the R package WGCNA^36^ with the transposed expression matrix as input. All expression features associated with each gene are integrated in Datafile 2.

### Heritability estimations

In a diallel scheme with no selfs (homozygous parental lines) and no reciprocals (half-diallel)^31^, the phenotype of the hybrid from crossing the *i* × *j* th parental lines can be expressed as:

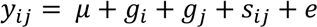

Where *µ* is the population mean, *gi* is the average contribution of all half siblings related to the *i* th parent, *gj* is the average contribution of all half siblings related to the *j* th parent (general combining abilities or GCA), and *sij* is the specific contribution of the *i* × *j* parental combination (specific combining abilities or SCA). Residual error is expressed as *e*.

We estimated these components with a linear mixed model using the lmer function from the R package lme4^37^. We excluded all homozygous hybrids and heterozygous hybrids with chromosome level allele imbalance (Table S2). In total, 258 hybrids were included in the model. We defined the GCA components as fixed effects and SCA as random effect. For each expression trait, we extracted the GCA and SCA estimates using the fixef and ranef functions, respectively. Error component is extracted as the residual value of the fitted model using the resid function. The variance components are estimated as:

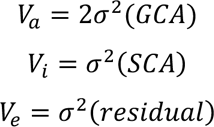

Where *Va* corresponds to the total additive variance, *Vi* is the non-additive variance due to interactions and dominance and *Ve* is the residual variance. The heritability is estimated as:

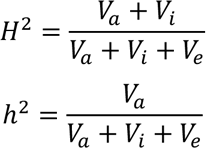

Where *H*^2^ is the broad-sense heritability and *h*^2^ is the narrow-sense heritability. The functions used for modeling and heritability calculations are available upon request as custom R scripts.

We applied an orthogonal strategy to estimate the narrow-sense heritability based on genome-wide kinship matrix (*h^2^g*). We first calculated the linkage disequilibrium using the SNP matrix and PLINK^38^. We excluded SNPs with strong linkage disequilibrium (r^2^ > 0.8) due to the strong population structure of the diallel panel. We calculated the weights of each variant using ldak with the –cut-weights and –calc-weights- all arguments and the default parameters^39^. All variants with non-zero weights were to generate a filtered vcf matrix of 5,493 SNPs that was then recoded with the plink -make-bed command. The filtered and recoded matrix was used to calculate the kinship between the individuals using the popkin function of the R package popkin with the default parameters. Finally, we used the hglm function of the hglm R package, with the default parameters, to calculate *h^2^g* from the kinship matrix and the tpm values. All heritability estimates are found in Table S4.

### Quantifying allele specific counts in the hybrids and the corresponding parental pairs

To infer the heterozygous sites in every hybrid, the sequencing reads of each parent from Peter *et al., 2018* were aligned to the R64_nucl reference genome using the *bwa mem* with the options *–M –t 8 –v 3* command and SNPs were inferred using *gatk HaplotypeCaller* ^40^. The vcf files of the parents of each cross were combined to generate a vcf file containing the heterozygous biallelic sites in the hybrid. We used ASEReadCounter^41^ to quantify the number of reads carrying each allele of the heterozygous biallelic sites. To avoid regions that would cause mapping problems, we calculated the mappability of 75bp segments along the R64_nucl reference genome using GenMap^42^ *genmap map –K 75 –E 2* and removed the sites with mappability values less than 1.

To quantify the read counts for the same discriminating sites in the parental samples, we calculated the depth of sequencing of the SNPs between the parent and the reference genome. The depth values were then scaled to the total number of reads for the hybrid, so that the depth values would be comparable across parents and the corresponding hybrid. We only considered sites with more than 30X coverage in the hybrids and more than 60X coverage in the sum of the parental pairs. In total, 1,864,327 sites were included for further analyses.

We calculated the chromosome-level allele balance across all hybrid samples as well as the corresponding parental pairs to identify systematic biases due to the presence of aneuploidy, loss-of-heterozygosity (LOH) or other large-scale chromosomal changes that could be frequent in *S. cerevisiae* cultures^43^. We plotted the allele balance (read counts for one parental line against the other at discriminating sites) and manually verified cases with inconsistent allele balances. In total, 258 hybrids that did not show such allele imbalance were included in the linear mixed model for heritability estimation. For allele-specific expression analysis, we further removed parent-hybrid trios where only the parental alleles were imbalanced, resulting in a set of 179 parent-hybrid trios. In total, these 179 parent-hybrid trios comprised 1,2M discriminating sites and are further analyzed.

### Allele-specific expression analyses across hybrid and their corresponding parental pair

We performed gene-level ASE analyses using the R package MBASED^44^. For each gene, all discriminating sites were considered as phased and were included to calculated the allelic change both in the hybrid and between the parental pair as 1-sample ASE test using 10^6^ simulations. Gene-level allele counts were approximated by averaging the counts at each site included in the test. The raw P-value were adjusted using the Benjamini-Hochberg method. For genes with more than 20 discriminating sites, only one site at each 100 bp window were sampled to remove redundancy due to overlapping reads. Significant 1-sample ASE test is considered when the adjusted P-value is less than 0.05 and the absolute log2 foldchange of the gene-level allele counts is higher than 1.

For genes where both the hybrid and parental 1-sample tests were significant, 2-sample tests were performed to identify significant ratio change between hybrid and parental ASE. We distinguished two scenarios. First, for cases where the signs of the log2 foldchanges in the parents and the hybrids are in the same direction, 2-sample tests were directly performed using the same sites and counts as the 1-sample tests. For cases there the signs were in opposite direction, the alleles in the hybrid were reverted before the test were performed. For 2-sample tests, 10^3^ simulations were performed and the P-values were adjusted using the Benjamini-Hochberg method. Significant 2-sample ASE test is considered when the adjusted P-value is lower than 0.05. Criteria for assigning gene-level regulation changes are in Figure 3D.

All ASE test results and the assigned regulation patterns are in Datafile 2.

### Gene-set enrichment analyses

Gene-set enrichment analyses (GSEA) were performed using the fgsea package in R^45^. Standard GO terms associated with biological processes (BP) were used^46^ with term size limits between 5 and 500. GSEA on different variance components were performed using rankings based on additive, non-additive and residual variance with 100,000 simulations. Results are found in Table S5. For GSEA on genes with different regulatory changes, the rankings were based on the number of cases for a given assigned pattern divided by the total number of cases characterized for that gene. The same number of simulations were performed as for the variance components enrichments. Results are found in Table S6.

### Assigning gene-level major regulatory patterns

We focused on *cis* only, *trans* only and compensatory assignments as they are the most major patterns observed across parent-hybrid trios. For a given gene, we calculated the number of regulatory patterns observed across trios. We assigned the gene-level regulatory pattern as “null” if the number of null patterns represent more than 95% of all characterized cases. For the remaining cases, we identified the most common regulatory change that is not null, then assigned that gene as mainly regulated by that pattern. All assignments are found in Table S7. We used the same criteria to assign the major regulatory groups for the within and between species allele pairs. Switches of the regulatory types are defined by comparing assigned types for between and within species pairs. All assignments are found in Table S8.

### Data availability

All RNA sequencing reads are available in the European Nucleotide Archive (ENA) under the accession number PRJEB64466.

The 1002 Yeast Genome website - http://1002genomes.u-strasbg.fr/files/ - (RNAseq) provides access to:

– Datafile S1: Datafile1_raw_data_20230713.tab
– Datafile S2: Datafile2_ase_sum_20230609.tab

## Acknowledgements

This work was supported by a National Institutes of Health (NIH) grant R01 (GM147040-01), a European Research Council (ERC) Consolidator grant (772505) to J.S and a French National Research Agency (ANR) young investigator grant (ANR-22-CE12-0023-01) to J. H. It is also part of Interdisciplinary Thematic Institutes (ITI) Integrative Molecular and Cellular Biology (IMCBio), as part of the ITI 2021-to-2028 program of the University of Strasbourg, CNRS, and Inserm, supported by IdEx Unistra (ANR-10-IDEX- 0002). J.S. is a Fellow of the University of Strasbourg Institute for Advanced Study (USIAS) and a member of the Institut Universitaire de France.

