## Supplemental Material for "Non-additive genetic components contribute significantly to population-wide gene expression variation"

for

**This document includes:**

Supplementary figures S1-S5

Supplementary figure legends

Descriptions for supplementary tables S1-S8

Online Datafile descriptions

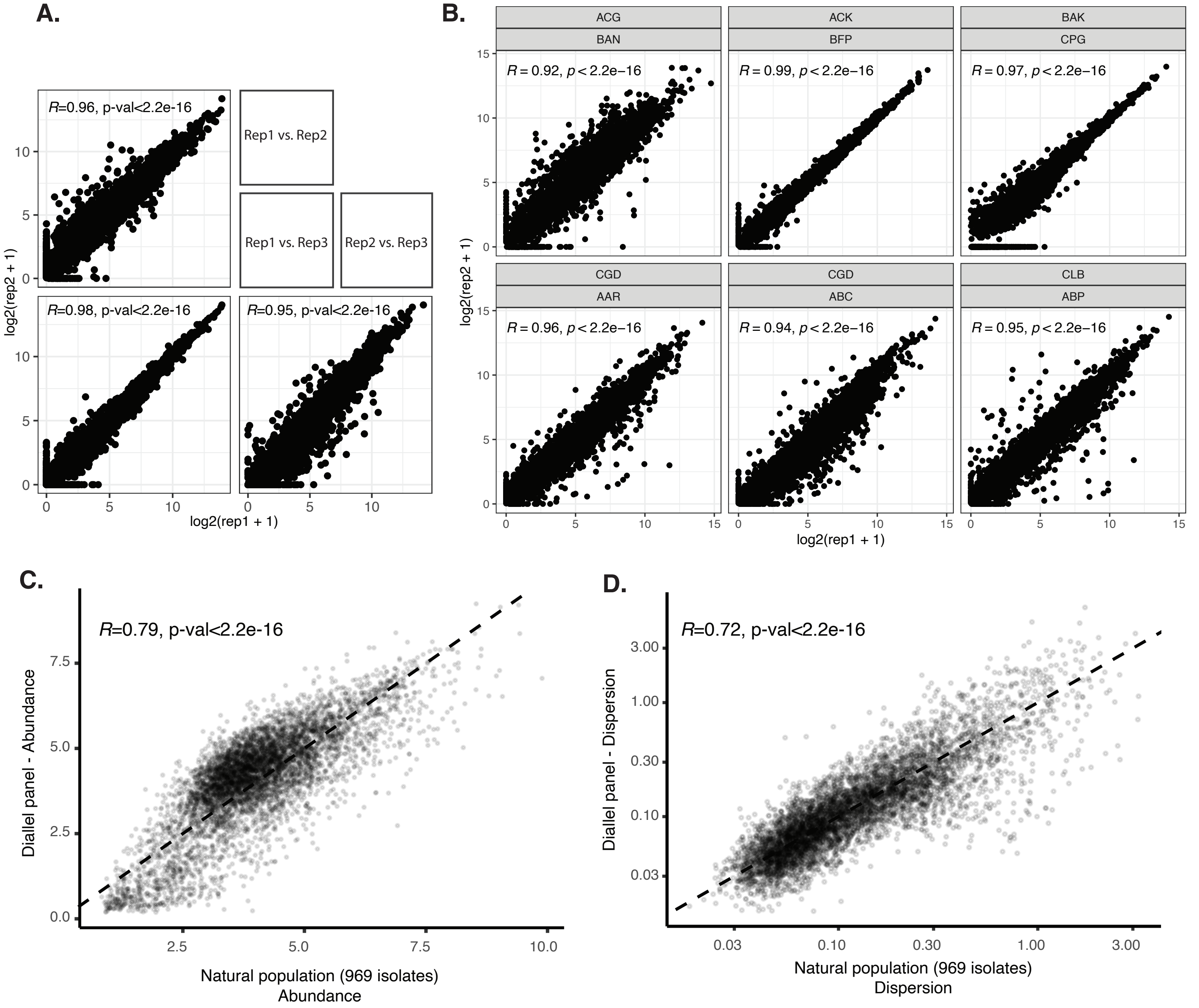


**Figure S1. Correlations of expression across samples and datasets.**
**A.** Pairwise correlations of three replicated samples of the homozygous diploid parental line AKE. Correlations are calculated based on the log2(tpm+1). **B.** Correlations between replicates of 6 heterozygous hybrids. Upper and lower label indicate the first and second parental lines, respectively. **C.** Correlation of the mean transcript abundance between data from a natural population of 969 isolates (x-axis) and the diallel panel (y-axis). Transcript abundance is calculated as log2(tpm+1). **D.** Correlation of the expression dispersion between data from a natural population of 969 isolates (x-axis) and the diallel panel (y-axis). Dispersion is calculated as the mean absolute deviation of transcript abundance. All correlations are based on Pearson's R.

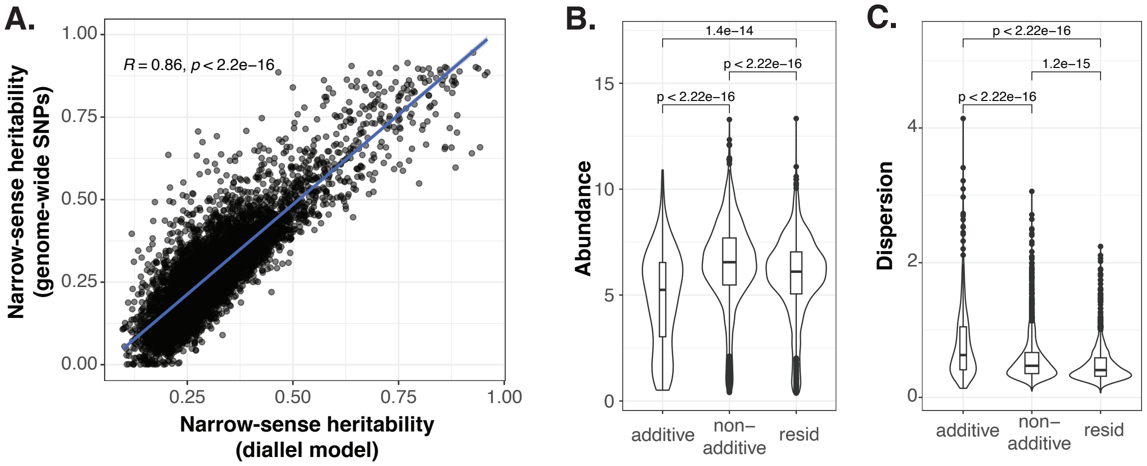


**Figure S2. Heritability and gene expression features for different variance components.** **A.** Correlation of narrow-sense heritability estimates based on the diallel model ($h^2$) (x-axis) and genome-wide SNP matrix ($h^2_g$) (y-axis). Correlation is based on Pearson's R. **B-C.** Distribution of the expression abundance (**B**) and dispersion (**C**) for genes that are mainly controlled by each of the variance components. P-values correspond to two-sided Wilcoxon tests.

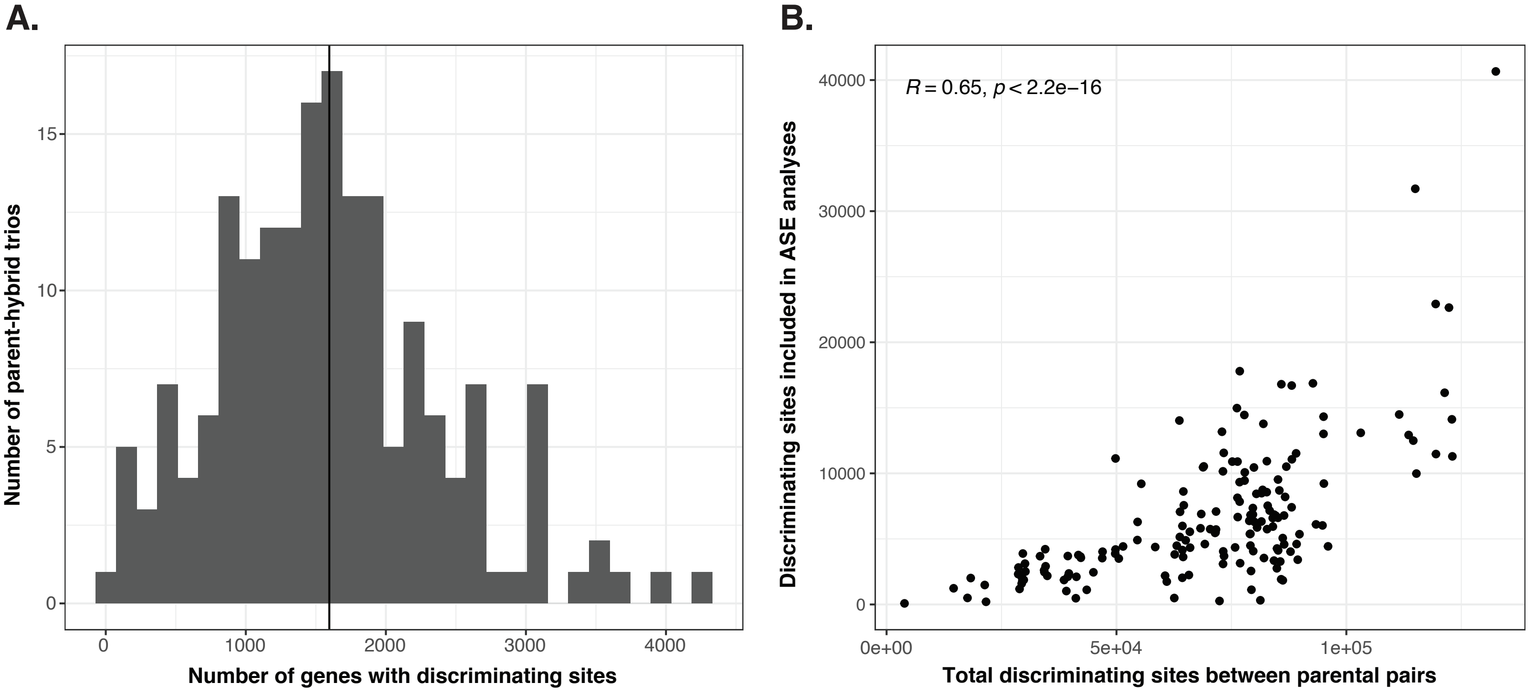


**Figure S3. Description of the allele-specific expression dataset.**
**A.** The distribution of the number of genes with at least one discriminating site across parent-hybrid trios (x-axis). The number of trios are indicated on the y-axis. The average gene number per trio is shown by the vertical black line. **B.** The number of discriminating sites included in the allele-specific expression analysis (y-axis) as a function of the total number of discriminating sites in each parental pair (x-axis). Each point represents a parental pair. Correlation coefficient based on Pearson's R.

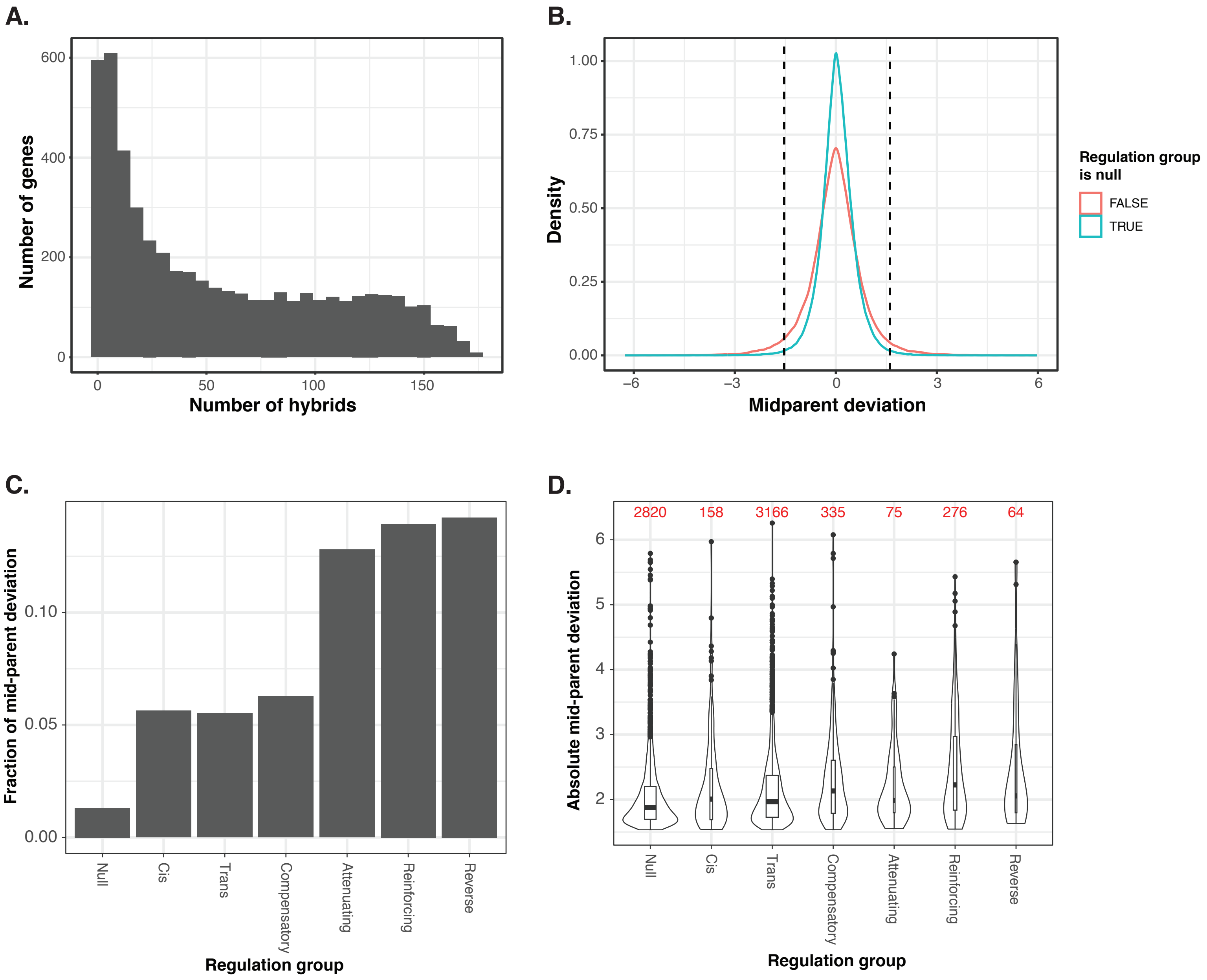


**Figure S4. Allele-specific expression analysis and mid-parent deviation.**
**A.** Distribution of the number of hybrids included for a given gene in the ASE analysis. **B.** Distribution of the midparent deviation for genes in the null regulation group (blue) and genes in other regulation groups (red). The dashed vertical lines represent the cutoff at top and bottom 2.5 percentiles. **C.** Fraction of mid-parent deviation cases for each of the 6 regulation groups and those in the null group. **D.** Box and violin plots depicting the absolute mid-parent deviation for the null and other regulation groups. The number in red mark the number of genes included in each group.

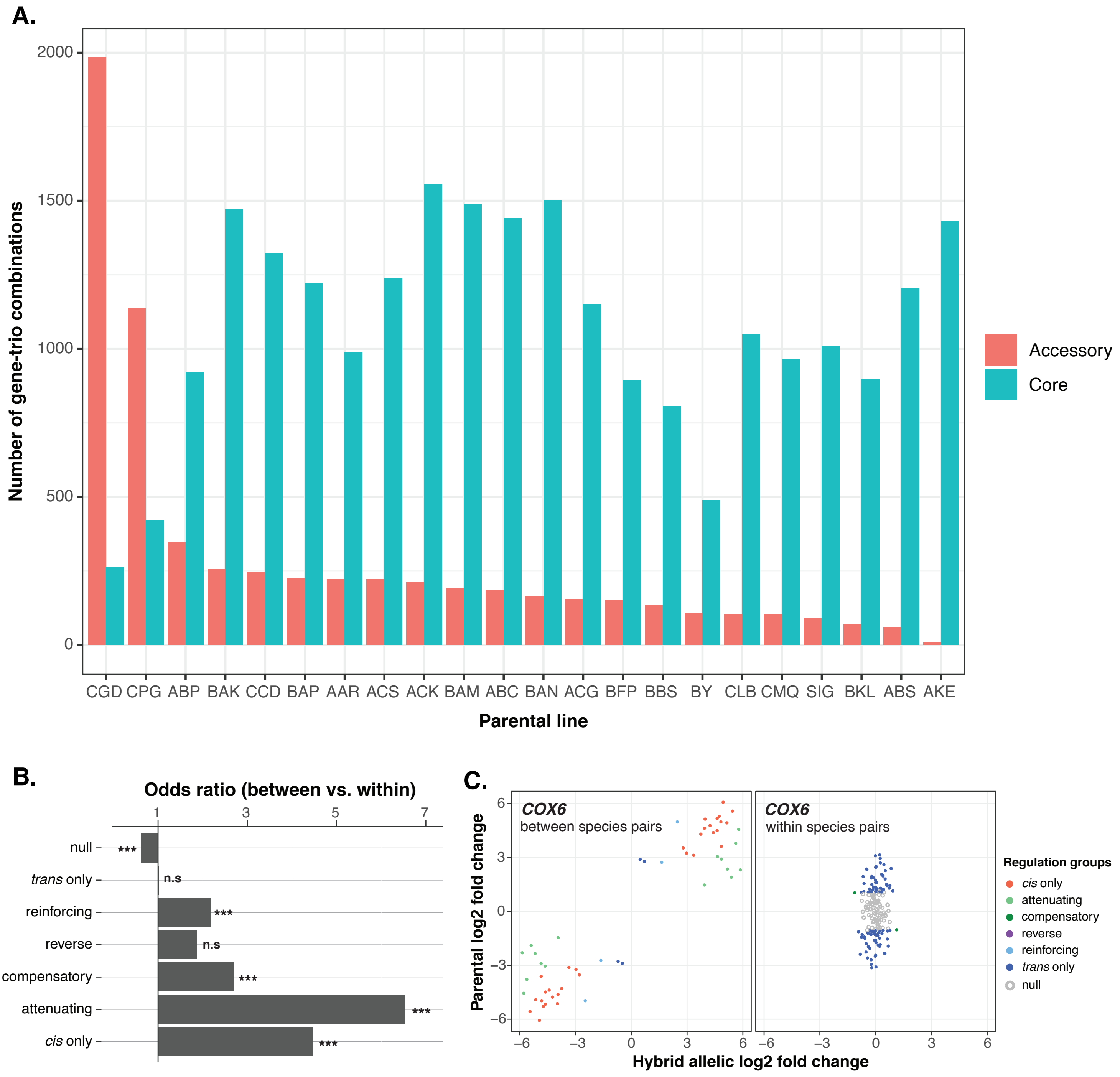


**Figure S5. Between and within species allele pairs and variation in regulatory patterns.**
**A.** The number of between (red) and within species (cyan) gene-trio combinations for each of the 26 parental line. **B.** Odds ratios (OR) comparing the regulatory patterns observed in between- vs. within-species pairs for the same set of 202 genes. OR and P-values are based on two-sided Fisher's exact tests. **C.** An example of regulation pattern switch between introgressed alleles (between species pairs) and native alleles (within species pairs). Regulatory patterns are color-coded. Log2 fold changes between alleles in the hybrid and between the parental lines at the same sites are indicated on x- and y-axes, respectively.

### Supplementary table descriptions

**Table S1-** Description of parental isolates included in this study.

**Table S2-** Description of hybrids included in this study.

**Table S3-** Description of genes included in this study.

**Table S4-** Heritability and genome-wide heritability estimates

**Table S5-** GSEA results across variance components

**Table S6-** GSEA results across gene expression regulatory variation

**Table S7-** Gene-level regulatory variation assignments

**Table S8-** Within and between species allelic regulatory variation assignments

### Online Datafile descriptions

**Datafile 1-** Normalized gene expression levels (TPM) and raw counts data across 331 samples

**Datafile 2-** Allele specific expression data across parent-hybrid trios
